# Programmed His bundle pacing - a novel maneuver for the diagnosis of His bundle capture

**DOI:** 10.1101/471268

**Authors:** Marek Jastrzębski, Paweł Moskal, Agnieszka Bednarek, Grzegorz Kiełbasa, Pugazhendhi Vijayaraman, Danuta Czarnecka

**Author notes:** Corresponding author: Assoc. Prof. Marek Jastrzębski, MD, PhD, First Department of Cardiology, Interventional Electrocardiology and Hypertension, Jagiellonian University, ul. Kopernika 17, 31-501 Kraków, Poland.

## Abstract

**Background:** During permanent non-selective (ns) His bundle (HB) pacing, it is crucial to confirm HB capture / exclude that only right ventricle (RV)-myocardial septal pacing is present. Because the effective refractory period (ERP) of the working myocardium is different than the ERP of the HB, we hypothesized that it should be possible to differentiate ns-HB capture from RV-myocardial capture using programmed extra-stimulus technique.

**Methods:** In consecutive patients during HB pacemaker implantation, programmed HB pacing was delivered from the screwed-in HB pacing lead. Premature beats were introduced at 10 ms steps during intrinsic rhythm and also after a drive train of 600 ms. The longest coupling interval that resulted in an abrupt change of QRS morphology was considered equal to ERP of HB or RV-myocardium.

**Results:** Programmed HB pacing was performed from 50 different sites in 32 patients. In 34/36 cases of ns-HB pacing, the RV-myocardial ERP was shorter than HB ERP (271.8±38 ms vs 353.0±30 ms, p < 0.0001). Programmed HB pacing using a drive train resulted in a typical abrupt change of paced QRS morphology: from ns-HB to RV-myocardial QRS (34/36 cases) or to selective HB QRS (2/36 cases). Programmed HB pacing delivered during supraventricular rhythm resulted in obtaining selective HB QRS in 20/34 and RV-myocardial QRS in 14/34 of the ns-HB cases. In RV-myocardial only pacing cases (“false ns-HB pacing”, n=14), such responses were not observed – the QRS morphology remained stable. Therefore, the PHB pacing correctly diagnosed all ns-HB cases and all RV-myocardial pacing cases.

**Conclusions:** A novel maneuver for the diagnosis of HB capture, based on the differences in ERP between HB and myocardium was formulated, assessed and found as diagnostically valuable. This method is unique in enabling to visualize selective HB QRS in patients with otherwise obligatory ns-HB pacing (RV-myocardial capture threshold < HB capture threshold).

**What this study adds:**

- Programmed His bundle pacing – a novel and straightforward method for unquestionable diagnosis of His bundle capture during non-selective pacing was developed and assessed.
- A method for visualization of selective HB capture QRS in patients with obligatory non-selective pacing (myocardial capture threshold < His bundle capture threshold) was discovered and physiology behind it explained.

## Introduction

The emergence of permanent His bundle (HB) pacing as a new, potentially alternative pacing option,^1–5^ poses new challenges in ECG interpretation and electrocardiographic assessment of capture during device implantation. In patients with HB pacing, it is imperative that HB capture is confirmed both at the time of implantation and during follow-up. The diagnosis of HB capture is mostly based on paced QRS morphology assessment. However, during non-selective (ns)-HB pacing, paced QRS complex is a fusion between right ventricular (RV) myocardial capture and HB capture with variable contributions of the HB / RV depolarization wavefronts to the fused QRS. This makes the diagnosis of HB capture / exclusion of pure RV-myocardial pacing not always straightforward. Currently, the differentiation between ns-HB and RV-myocardial pacing is predominantly based on differences in capture thresholds between the HB and the RV myocardium. By increasing and decreasing pacing output, sudden changes of QRS morphology is observed reflecting HB/RV-myocardium capture/loss of capture.^6^ This method has limitations and can fail when 1.) the capture thresholds are similar, 2.) the change in QRS morphology is small/ambiguous, or 3.) the change in QRS morphology has a different cause than HB/RV capture/loss of capture. In clinical practice, it is quite challenging to determine if the HB is really paced or just pure RV-myocardial pacing with a relatively narrow, ‘septal’ QRS complex present.^7^

The aim of this study is to assess a novel method for confirmation of HB capture during non-selective pacing. We hypothesized that since effective refractory period (ERP) of HB is different from the ERP of the RV myocardium, it should be possible to differentiate pure myocardial capture from ns-HB capture using programmed extra-stimulus testing. When the RV-myocardial ERP is shorter than the HB ERP, the first extra-stimulus delivered at a coupling interval shorter than the HB ERP should result in a sudden QRS widening revealing QRS morphology of pure myocardial capture; this finding would be diagnostic of non-selective HB pacing. In cases where the RV-myocardial ERP > HB ERP, the first extrastimulus with a coupling interval shorter that RV-myocardial ERP should be followed by isoelectric interval and then selective HB paced QRS complex. Such a response would also be diagnostic of ns HB capture. If no QRS morphology change is observed throughout the whole coupling interval range despite reaching refractoriness / complete loss of capture, this would be indicative that only pure RV-myocardial capture was present.

## Methods

In consecutive patients who underwent permanent HB pacemaker implantation, programmed HB pacing was performed and analyzed with the use of an electrophysiology system (Lab System Pro, Boston Scientific / Bard, USA). Pacing was delivered from the already deployed HB pacing lead (active helix, screw-in lead, model 3830, Medtronic, USA) with output set at 2 times the HB / RV capture threshold to ensure that both RV and HB were simultaneously captured (non-selective HB pacing, as recently defined).^6^ A change in QRS morphology during decrease / increase in the pacing output served as the gold standard to identify pure RV-myocardial paced QRS morphology and non-selective HB paced QRS morphology. Premature beats were introduced during the intrinsic rhythm and also after an 8-beat basic drive train of 600 ms. The coupling interval was decreased at 10 ms steps, starting from 450 ms, until complete loss of capture. If the 3830 lead was deployed at a different sites in the same patient – programmed pacing was repeated from the new site.

QRS morphologies obtained during programmed pacing were compared with the QRS morphologies obtained during differential pacing output technique and analyzed according to the hypothesized diagnostic principle delineated in the introduction section.

The effective refractory period was defined as the longest coupling interval between the last stimulus of the drive train and the premature stimulus that failed to depolarize the tissue. Relative refractory period was defined as the longest coupling interval that resulted in prolonged conduction as evidenced by QRS prolongation or stimulus-QRS interval prolongation.

All patients gave written informed consent for participation in this study and the Institutional Bioethical Committee approved the study protocol.

## Results

Consecutive patients (n=32), who underwent permanent HB pacemaker implantation were studied; clinical characteristics of these patients are presented in Table 1. Programmed HB pacing was performed 96 times from 50 different sites where the HB pacing lead was screwed-in during the procedure; 46 times during intrinsic supraventricular rhythm and 50 times with the use of an 8-beat basic drive train of 600 ms. In vast majority of the studied patients (30/32), the RV-myocardial refractory period was shorter than the HB refractory period. Consequently, for the whole group, the average RV-myocardial refractory period was significantly shorter than the HB refractory period: 271.8±38 ms vs 353.0±30 ms, p < 0.0001 and 306.4±37 ms vs 383.7±54 ms, p < 0.0001 when assessed with a drive train or with extrastimuli delivered during supraventricular rhythm, respectively. In all cases of ns-HB pacing (n = 36) when refractoriness was assessed with a drive train, there was a typical abrupt change of QRS morphology (QRS duration prolongation, rounding of R wave peak, appearance of notches, etc. – see Figure 1 and Supplementary Figure 1) occurring at a coupling interval range close to the expected refractoriness of the HB. The RV-myocardial paced QRS morphology was then maintained until the complete loss of capture at the RV myocardial refractory period. In all cases, this broader QRS morphology was identical to the RV myocardial QRS morphology obtained with differential output pacing maneuver in a particular patient (compare Supplementary Figure 1 and Supplementary Figure 2). This typical QRS morphology change was in some cases preceded by 1-2 slightly broader ns-HB QRS complexes likely due to the decremental conduction in the HB during the relative refractory period and hence a smaller contribution of the HB capture to the fused QRS complex – see Supplementary Figure 1.

**Figure 1.**
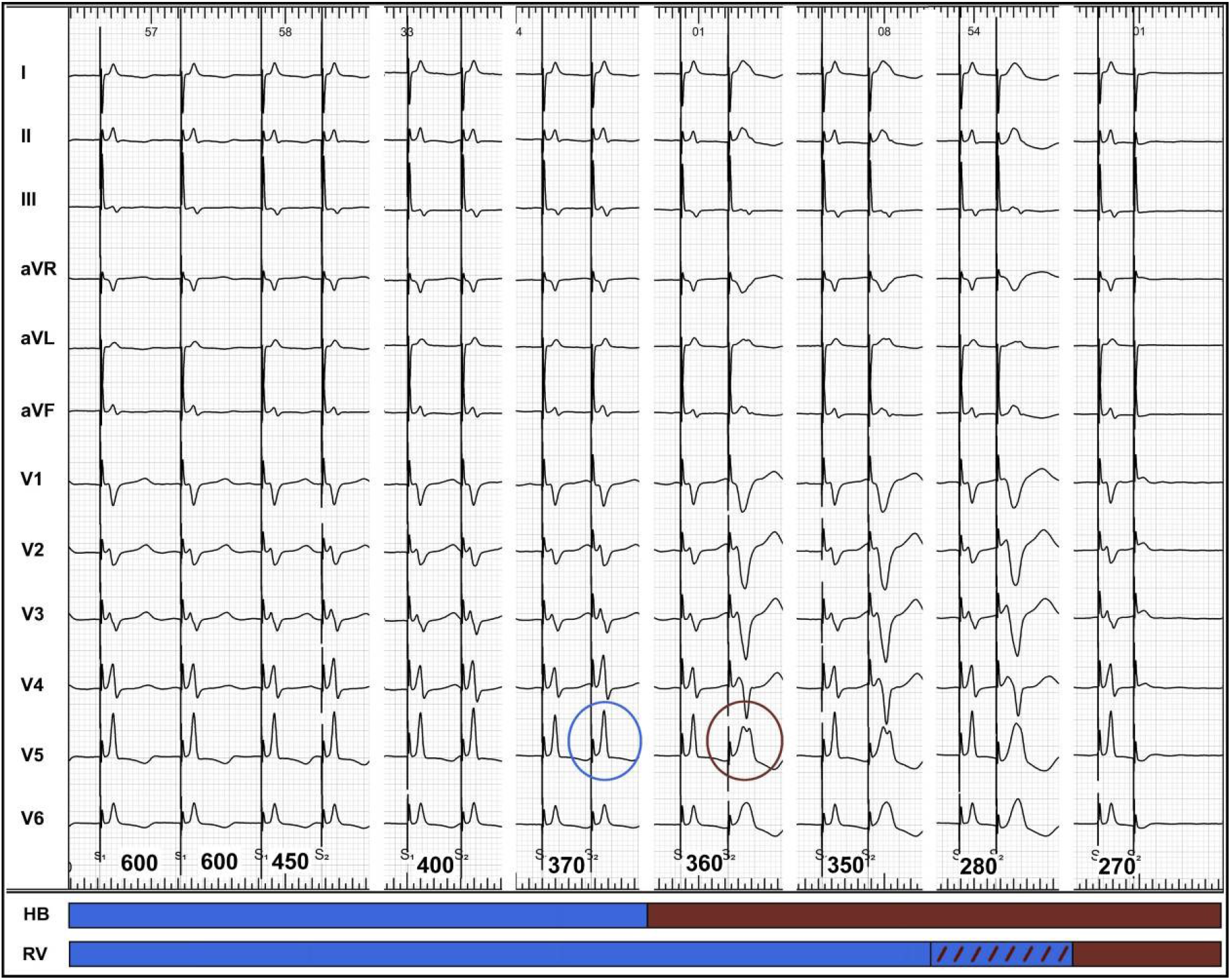
Programmed HB pacing: premature extra-stimuli are delivered after a drive train of 600 ms at progressively shorter coupling intervals. At coupling interval of 370 ns-HB, QRS morphology is still present (blue circle) while at a 10 ms shorter coupling interval of 360 ms, HB is found refractory and pure RV-myocardial morphology is unmasked (burgundy circle). At coupling intervals of 450 – 370 ms, only ns-HB QRS morphology is present (HB effective refractory period of 360 ms). At coupling intervals of 360-280 ms, only RV-myocardial QRS morphology is present (RV myocardial effective refractory period of 270 ms). During relative refractory period of the RV myocardium (280-270ms), some further QRS prolongation can be observed. The blue bar corresponds to HB / RV capture (excitable period), the dashed bar to capture with decremental conduction (relative refractory period), and the burgundy bar to loss of capture (effective refractory period).

**Table 1.** Basic clinical characteristics of the studied group (n = 32)

|  |  |
| --- | --- |
| Age [years] | 75.1±12.6 |
| Male gender | 21 (65.6%) |
| Pacing indication: |  |
| • Sick sinus syndrome | 3 (9.4%) |
| • Atrioventricular block | 8 (25.0%) |
| • Atrial fibrillation with bradycardia | 13 (40.6%) |
| • Heart failure | 8 (25.0%) |
| Procedure result: |  |
| • HB pacing | 20 (62.5%) |
| • Myocardial / deep septal pacing | 12 (37.5%) |
| Comorbidities: |  |
| • Heart failure | 15 (46.9%) |
| • Coronary heart disease | 11 (34.4%) |
| • Diabetes mellitus | 13 (40.6%) |
| • Hypertension | 25 (78.1%) |
| • Severe valvular disease | 4 (12.5%) |
| LV ejection fraction [%] | 49.3±11.2 |
| Native QRS duration [ms] | 123±25.4 |
| HV interval [ms] | 50.2±9.6 |

HB – His bundle, ns – non-selective, LV – left ventricular, HV – His-ventricle

In two patients, instead of RV-myocardial QRS a selective HB QRS appeared with short coupled extrastimuli (RV-myocardial ERP > HB ERP). In stark contrast to the response observed in ns-HB pacing cases, in all cases of pure RV myocardial capture (‘false ns-HB pacing’, n = 14) there was no change of QRS morphology throughout the whole coupling interval range – see Figure 2.

**Figure 2.**
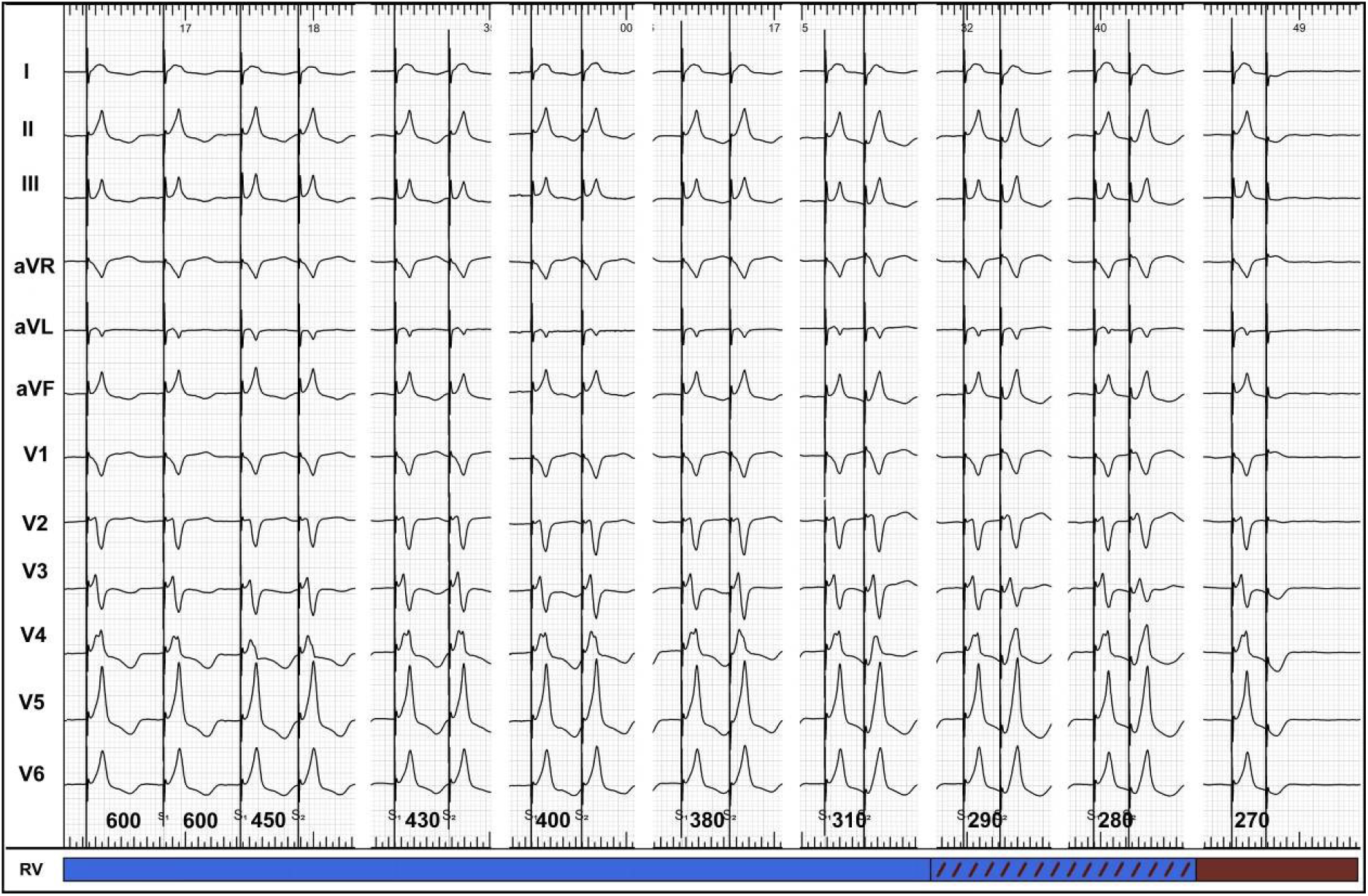
Programmed HB pacing during RV-myocardial capture only (‘false ns-HB pacing’): stable QRS morphology is present until RV-myocardial effective refractory period and loss of capture that occurs within the usual coupling interval values (280-250 ms).

Surprisingly, extra-stimuli delivered during native supraventricular rhythm resulted in the majority of cases of ns-HB pacing (20 out of 34) in a ‘reversed response’. Short-coupled extrastimuli brought out pure selective HB capture QRS morphology (Figure 2) rather than pure RV-myocardial morphology. This phenomenon of sudden loss of RV-myocardial capture was present despite the fact that the RV refractory time was shorter than the HB refractory time and despite pacing with output 2x higher than the RV-myocardial capture threshold. In 11/34 cases, there was no ‘reversed response’ but the same response as during programmed pacing with the drive train, i.e. appearance of RV-myocardial QRS morphology when HB refractoriness was met and in 3/34 cases ns-HB QRS morphology was maintained until loss of capture (simultaneous loss of capture of the HB and RV-myocardium).

In summary, programmed HB pacing provided a diagnostically correct response in every studied patient, both when premature beats were introduced during intrinsic rhythm and when an 8-beat basic drive train of 600 ms was used. Representative examples of responses during programmed HB pacing are presented in Figures 1–3 and Supplementary Figures 1 and 3.

**Figure 3.**
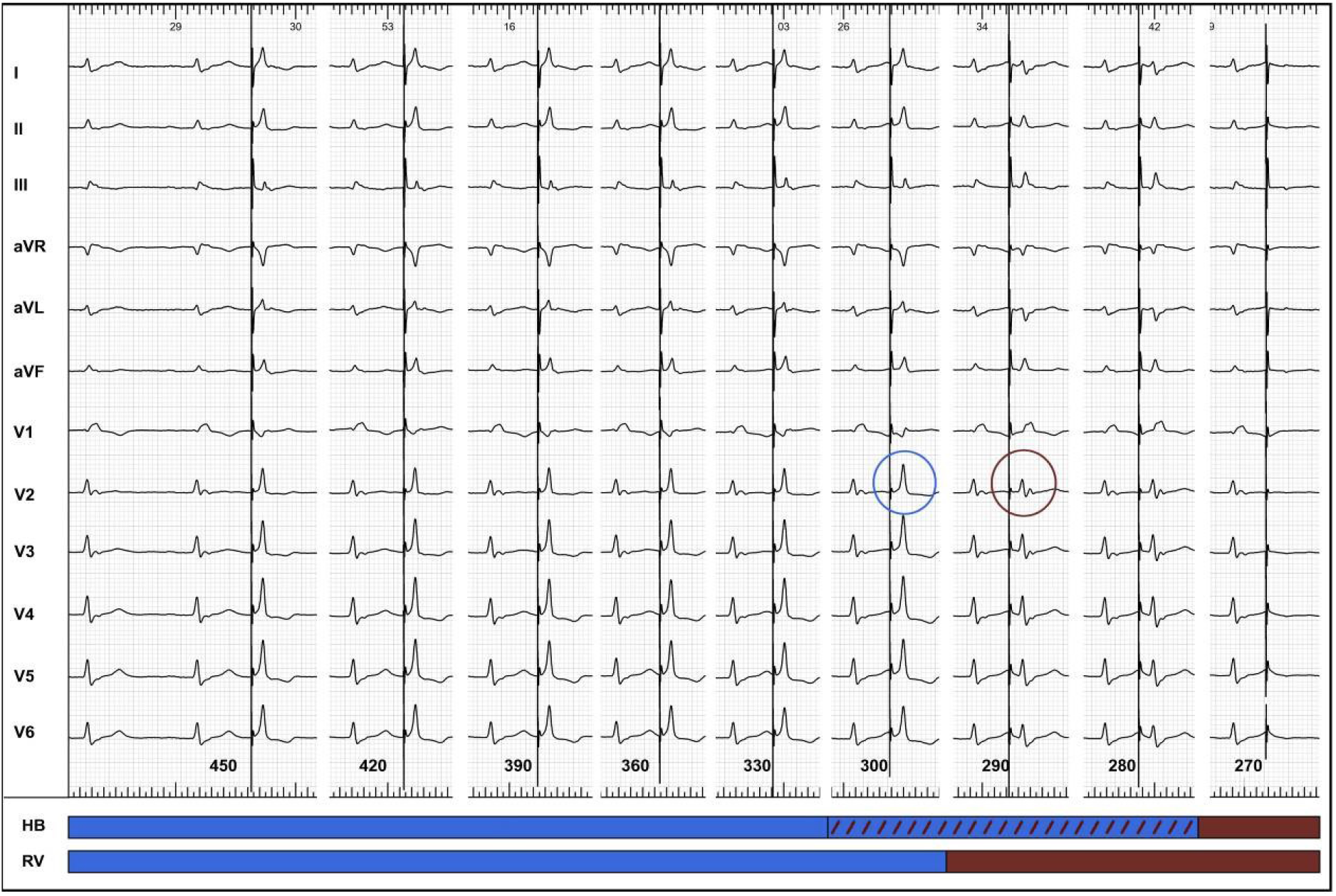
Programmed HB pacing during native supraventricular rhythm. Blue and burgundy circles denote the sudden change of QRS morphology from ns-HB QRS morphology to selective-HB QRS morphology. At coupling intervals of 450 – 300 ms, only ns-HB QRS morphology is present. At coupling intervals of 290-270 ms, only selective-HB QRS morphology is present. This reflects RV refractory period of 290 ms and HB refractory period of 370 ms (270 ms + 100 ms) – for explanation, see text and Figure 4. Note that at the coupling interval of 300 ms, there is some QRS prolongation, most likely due to the diminished contribution of the HB depolarization wavefront to the fused ns-HB QRS. This is most likely caused by the encroachment on the relative refractory period of the HB and hence the HV interval prolongation. Long stimulus-V interval present with the coupling interval of 290 ms and further prolongation with subsequent extra-stimulus (280 ms) supports such an explanation. The blue bar corresponds to HB / RV capture, the dashed bar to capture with decremental conduction (relative refractory period), and the burgundy bar to loss of capture (effective refractory period).

## Discussion

The major finding of the current study is that the programmed HB pacing maneuver is able to reliably differentiate non-selective HB capture from RV-myocardial pacing. Diagnostically correct response was observed in all cases of ns-HB pacing, because ample difference in refractory times between HB and RV myocardium was present in every studied patient. Classic studies on the refractoriness of the human heart provide concordant data with regard to the difference in refractory times between the HB and RV myocardium.^8^ It seems that this electrophysiological characteristics of the heart can be relied upon – for diagnostic purpose – in patients with permanent HB pacing.

The second important and novel finding of this study is that the programmed HB pacing method is uniquely capable of visualizing selective HB QRS morphology in cases where this seemed impossible, i.e. when RV-myocardial capture threshold is lower than the HB capture threshold leading to obligatory RV capture during HB pacing. We believe that this interesting phenomenon of bringing out selective HB QRS complex can be explained by the altered activation sequence when premature stimulus is delivered after slower native supraventricular rhythm vs. after the faster basic drive train paced QRS. This is explained graphically on Figure 4. Briefly, when an extra-stimulus is delivered during supraventricular rhythm, the RV coupling interval is shorter than HB coupling interval. This is because HB is pre-excited in relation to the local ventricular myocardium near the HB pacing lead. The HB activation starts approximately 100 ms before the RV myocardium because of the sum of HV interval (50 ms) and the time necessary for the depolarization to reach the most basal part of the interventricular septum near the HB (50 ms). The average difference in refractory periods between HB and RV myocardium is about 80 ms, therefore, in most patients, this will result in a 20 ms longer excitable period of the HB than RV leading to selective HB QRS complexes at the two last coupling intervals before the ERP of the HB is reached. In contrast, during basic drive train both the HB and the local RV myocardium are depolarized simultaneously and the refractory periods of both structures begin simultaneously rather than sequentially, and selective HB capture is not observed since HB effective refractory period is almost always longer than RV effective refractory period.

**Figure 4.**
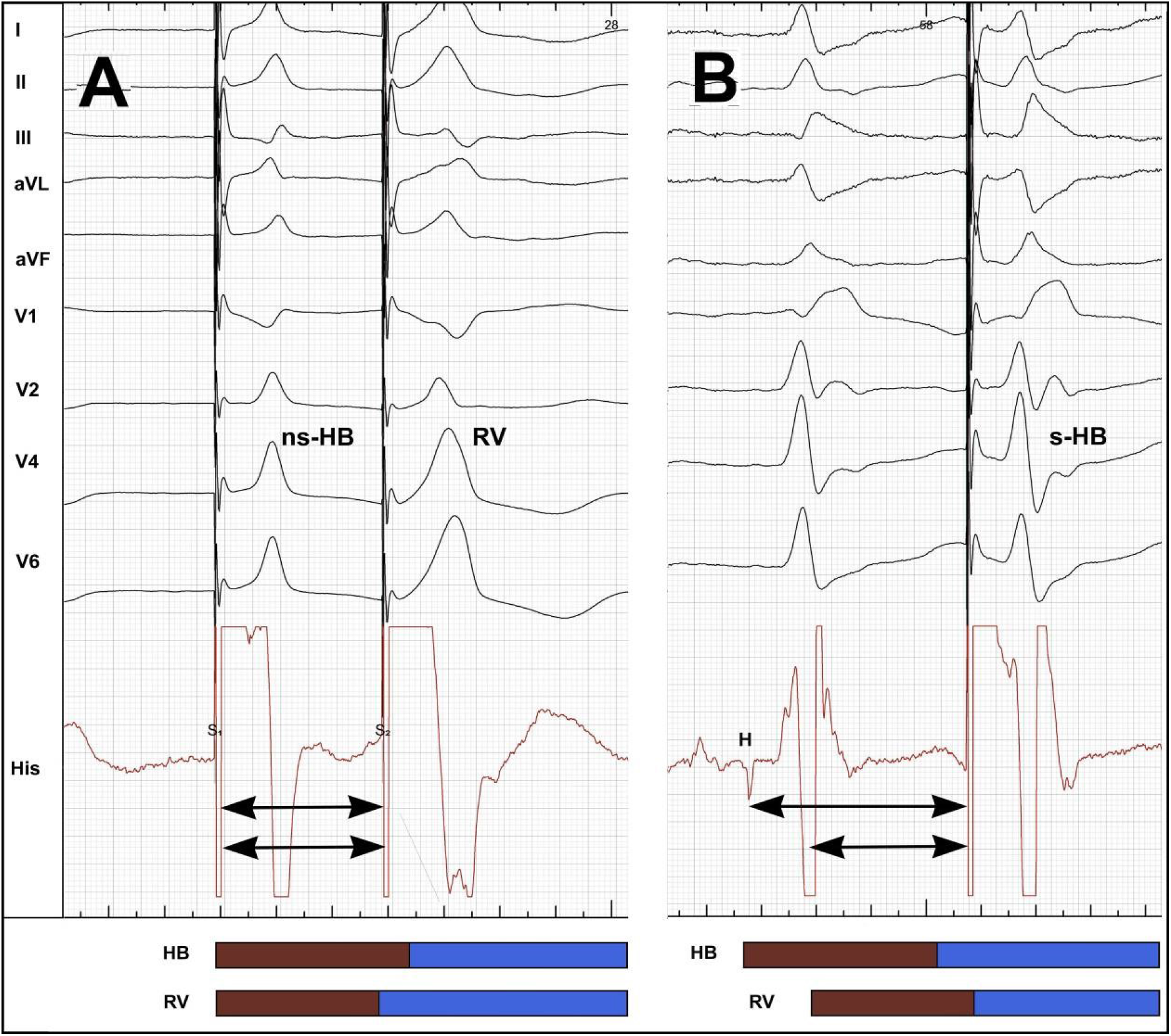
A “reversed response” during programmed HB pacing enables to visualize both components of the fused ns-HB QRS complex. In the same patient with a ns-HB pacing, an extra-stimulus is delivered at a coupling interval of 300 ms, first after a drive train (panel A) and then during a supraventricular rhythm (panel B). After a 600 ms drive train, the extrastimulus results in a sudden QRS broadening in comparison to the preceding ns-HB QRS because of RV-myocardial capture only. During the supraventricular rhythm, an extrastimulus with identical coupling interval unexpectedly resulted in a selective HB capture and QRS similar to the native supraventricular rhythm. This can be explained by a different activation sequence. During supraventricular rhythm, HB is activated 100 ms before the ventricular myocardium near the HB pacing lead. Consequently, despite a longer refractory period (burgundy bar), HB is excitable (blue bar), while RV myocardium is not. During the 600ms drive train of ns-HB pacing, both the HB and the local RV myocardium are depolarized at the same time and HB is not excitable due to the longer effective refractory period. Arrows mark the extent of the true coupling intervals for the HB and the RV. HB – His bundle, s-HB – selective HB, ns-HB – non-selective HB, His – endocardial signals from the screwed-in HB pacing lead, H – HB potential, RV – right ventricle

The two tested methods of programmed HB pacing seem complementary to visualize both components of the fused ns-HB QRS complex. While both methods provided diagnostic response in our cohort, these responses were different. Extra-stimuli delivered after a drive train best expose the difference in refractory periods between HB and RV myocardium and unmask pure RV-myocardial QRS. However, the responses observed when extra-stimuli are delivered during supraventricular rhythm seemed more diagnostically clear-cut as the selective HB QRS is diagnostically unmistakable.

### Clinical translation

In our experience, during the last 4 years and over 240 cases of permanent HB pacing device implantation, we have encountered several situations where the paced QRS morphology was ambiguous and either there was no change of QRS morphology with differential output method or the change was inconclusive and we were searching for an alternative diagnostic option. Without such a method, the operator faced a dilemma: should the lead be repositioned or was the acute endpoint of HB pacemaker implantation procedure already achieved? This problem is even more pronounced during follow-up when pacing from other sites for comparison is unavailable and the decision to schedule the patient for HB lead revision is much more serious. We believe that the programmed HB pacing maneuver and the principle behind it are the needed solution to this problem. We found that programmed HB pacing is useful for making the diagnosis of HB capture in ambiguous cases and for providing additional evidence of HB capture in the remaining obligatory ns-HB pacing cases by visualizing selective capture QRS. It is worth to note that the diagnostic value of a selective HB paced QRS exceeds that of any ns-HB paced QRS even when QRS morphology change is observed with differential pacing output technique. Programmed HB pacing assures the operator that the acute endpoint of the procedure was unquestionably achieved.

Programmed HB pacing method is not limited to use in the electrophysiology laboratory only, as it can be delivered from an implanted pacemaker as well, using the ‘non-invasive programmed stimulation’ option, that is available in most pacemakers. Perhaps it is not necessary to perform pacing with the whole coupling interval range. It seems that it is adequate to introduce a single extra-stimulus at a single coupling interval of 300 ms to obtain a straightforward diagnostic response (Supplementary Figure 3) in the majority of cases. We found that at this coupling interval HB is almost always refractory, while myocardium is almost always excitable. Moreover, the difference in ERP between the HB and RV myocardium can be exploited in a simplified manner. It is enough to temporarily program HB pacemaker to an asynchronous VOO mode with the pacing rate slightly slower than the native ventricular rate and to observe the paced QRS morphology behavior. Asynchronous pacing results in scanning of diastole with pacing stimuli, and it takes a short time to observe a diagnostic response: either broader RV-myocardial or narrower selective HB QRS complexes appear with short coupled stimuli (Figures 5 and 6).

**Figure 5.**
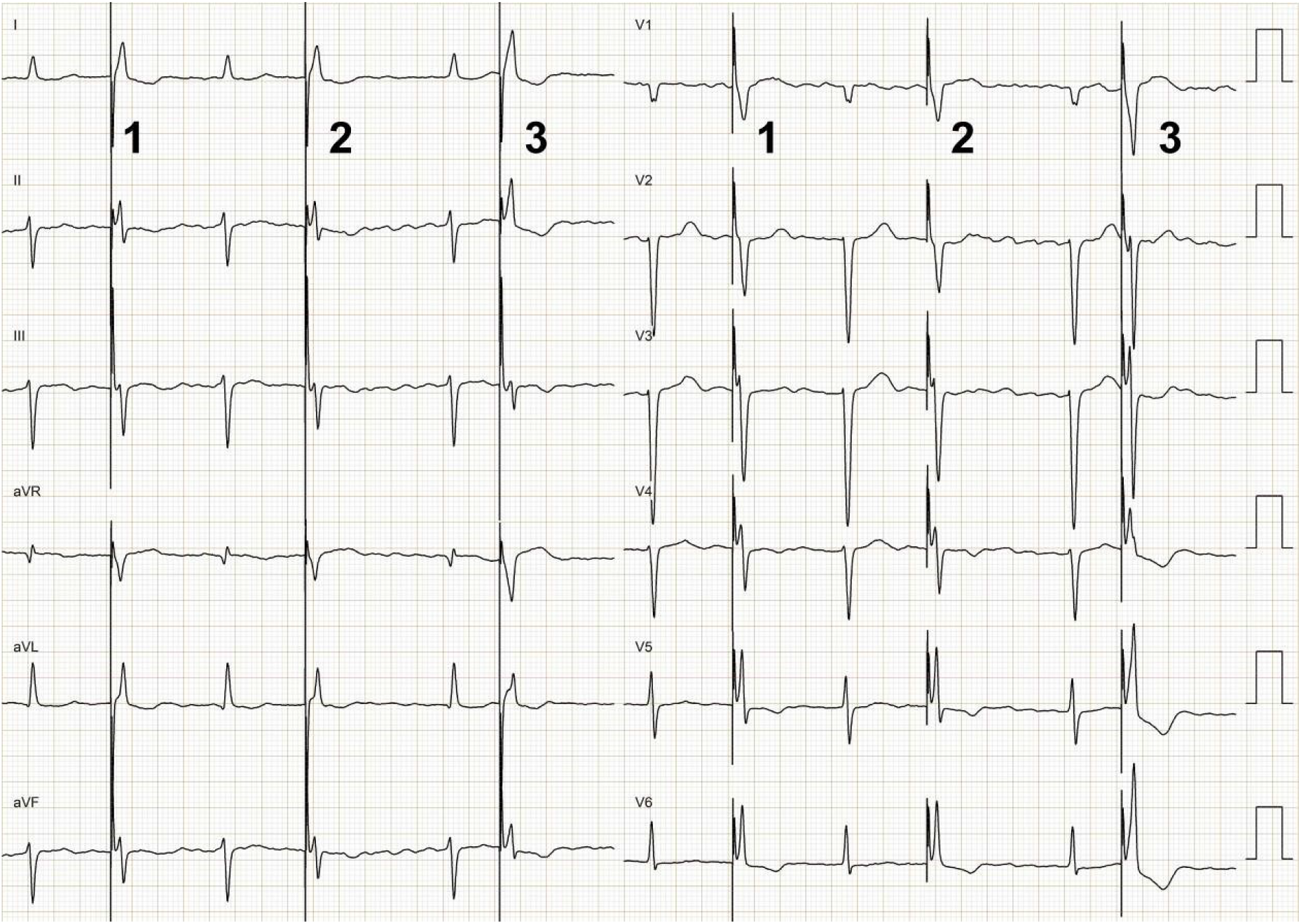
Asynchronous ns-HB pacing in VOO mode results in scanning of diastole with pacing stimuli. The first two stimuli result in ns-HB capture while the third occurs when the HB is refractory and depolarizes only the RV myocardium (note QRS axis change and QRS broadening) – thus providing a diagnostic response.

**Figure 6.**
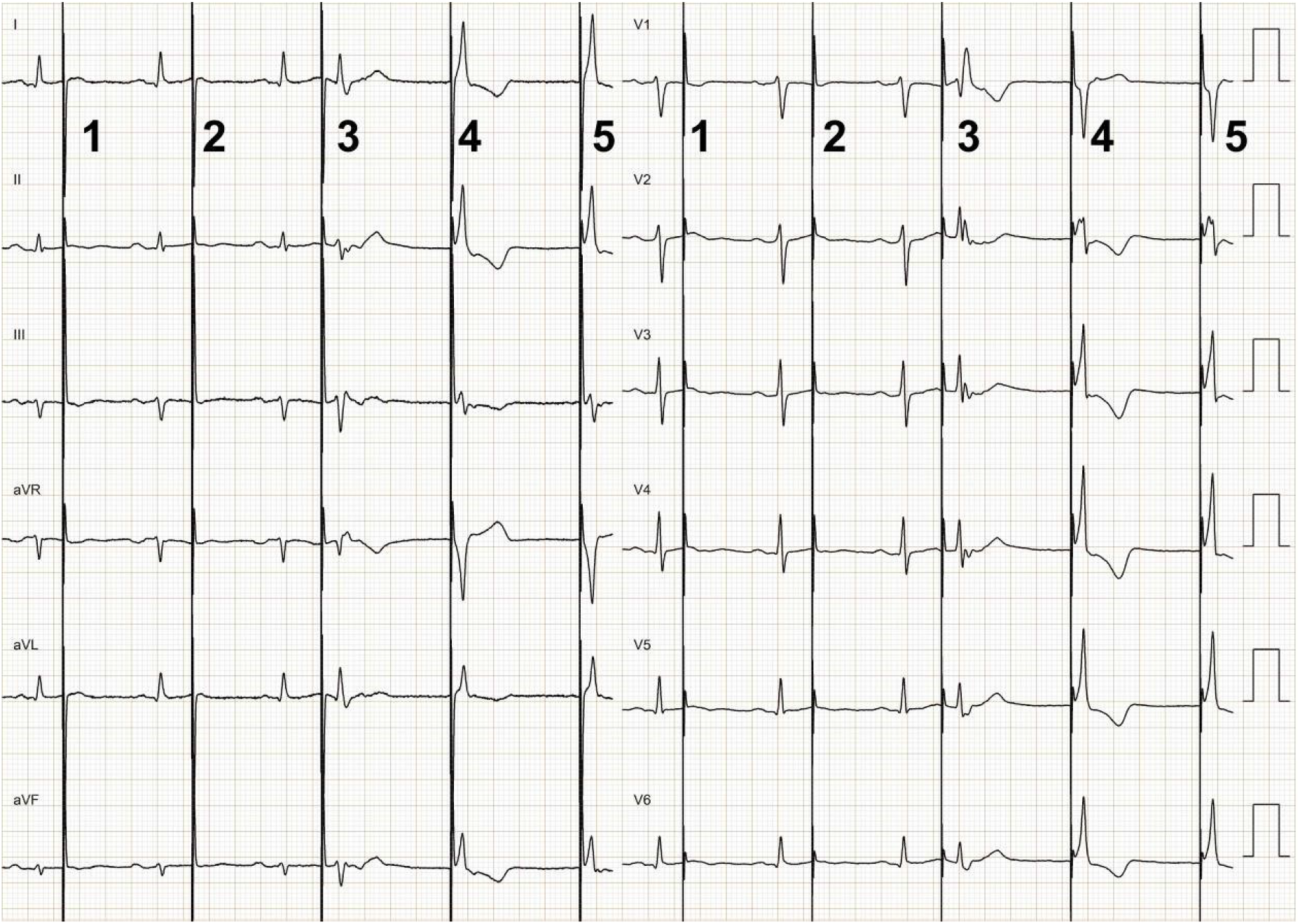
Asynchronous ns-HB pacing in VOO mode results in scanning of diastole with pacing stimuli. The first two stimuli fall on the effective refractory period of the HB and RV myocardium and result in non-capture, while the third stimulus occurs when the RV myocardium is refractory and propagates only via HB resulting in the short isoelectric interval and supraventricular QRS morphology with right bundle branch aberration (Ashman phenomenon). Stimuli 5 and 6 result in ns-HB capture since with a longer coupling, both the HB and RV myocardium are excitable.

### Other methods to differentiate ns-HB capture from septal myocardial capture

We are not aware of any validated method for differentiation between ns-HB capture and RV myocardial capture other than assessing changes in retrograde conduction.^7,9^ Although we did not assess VA conduction times in this study, this method has several limitations. First, in the most common scenario when an additional method is needed, i.e. no QRS morphology change with differential output due to similar capture thresholds, there also would be no change in VA conduction time. Second, in our experience, the majority of patients undergoing HB pacemaker implantation either have permanent atrial fibrillation or no 1:1 VA conduction, which precludes application of this method, as well.

### Limitations of the study

A relatively small number of patients were included, however, the observed responses were absolutely repeatable and consistent in the whole studied group, and the physiological explanation behind the observed responses is theoretically sound. We believe that the studied cohort was large enough to provide the necessary ‘proof of concept’ for the programmed HB pacing maneuver while ‘real life’ clinical usefulness needs to be shown in a bigger cohort. Based on our observations, the incidence of identical RV and HB capture thresholds in patients with ns-HB pacing is less than 10%. This was not implicitly assessed in this cohort.

Potential pitfall in interpretation of responses to programmed HB pacing that became apparent during the current study is the presence of QRS widening and HV interval prolongation when the RV and the HB relative refractory periods are encroached upon with extra-stimuli, respectively. These phenomena can be appreciated when analyzing QRS complexes captured during relative refractory periods in Figures 2–3 and Supplementary Figure 1. Prolongation of QRS complexes during relative RV myocardial refractory period should not be mistaken as prolongation due to the loss of HB capture. Features than can help in distinguishing between these two phenomena are as follows: 1.) QRS prolongation related to the relative refractory period is usually more gradual, subtle and present just before the effective refractory period i.e. 10 – 30 ms before the loss of capture; 2.) it is present at coupling intervals below the typical HB refractory times, i.e. < 300 ms, usually close to 260 ms 3.) an already broad and notched QRS, usually > 150 ms, changes into even more broader QRS.

The prolongation of the HV interval that occurs at coupling intervals just before the coupling interval with the loss of HB capture is responsible for QRS morphology change occasionally, somewhat visually less evident than expected (compare Figure 1 vs Supplementary Figure 1). HV interval prolongation results in the diminishing contribution of the HB capture to the fused QRS. Such a gradual QRS broadening at coupling intervals just before the coupling interval with complete loss of HB capture might causes difficulties in determining at exactly what coupling interval HB capture was lost. Analysis of all 12-ECG leads overcomes this problem, as in some leads change in QRS morphology/axis is more abrupt/evident (note lead III in Supplementary Figure 1 and 2) than in others.

Programmed ventricular stimulation is time-consuming and has the potential to induce malignant ventricular arrhythmias and this might be seen as a practical limitation of the current maneuver. However, programmed stimulation is considered safe when performed in an electrophysiology laboratory and induction of ventricular tachyarrhythmia with a single extra-stimulus is “extremely uncommon”.^8^ During our study, we did not observe induction of any ventricular arrhythmias (couplets or non-sustained ventricular tachycardia). Perhaps programmed HB pacing likely is less arrhythmogenic than programmed ventricular stimulation.

The same applies to the proposed temporary asynchronous ventricular pacing – it might be seen as proarrhythmic. However, asynchronous ventricular pacing is an officially recommended method used during pacemaker follow-up and even for remote telephone monitoring – apparently without negative consequences.

## Conclusions

A novel maneuver for the diagnosis of HB capture, based on the differences in ERP between myocardium and HB was formulated, assessed and explained. We believe that this diagnostic method increases knowledge of HB electrophysiology, provides a diagnostic solution in ambiguous paced QRS morphologies, and contributes to a more rigorous definition of the procedure endpoint. A larger study is necessary to fully evaluate the diagnostic value and clinical utility of this maneuver.

## Disclosures

P. Vijayaraman:

Research, Fellowship support, Speaker, Consultant – Medtronic

Consultant – Abbott, Biotronik and Boston Scientific

Speaker – Merritt medical.

## Supplementary Figures

**Supplementary Figure 1.**
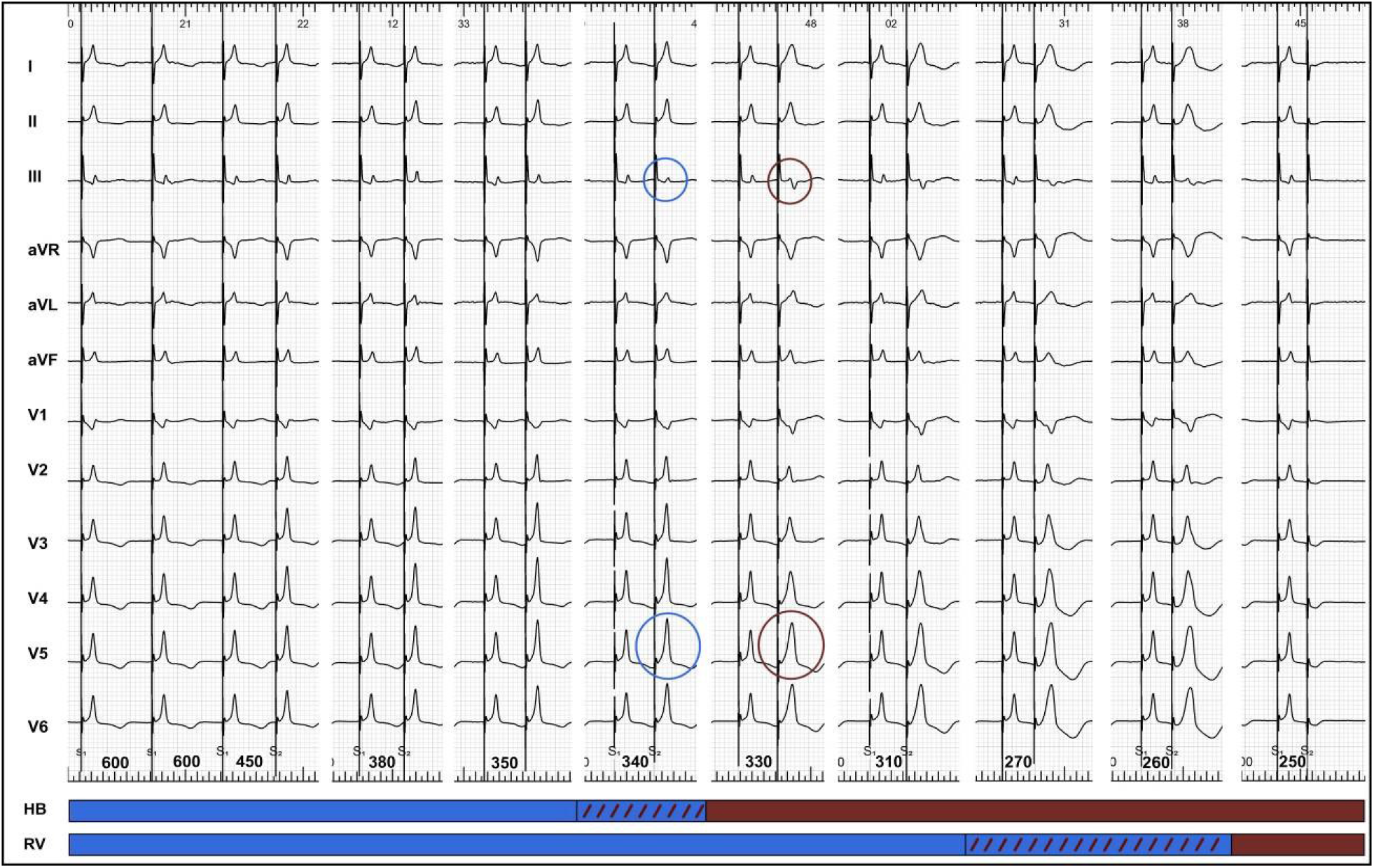
Programmed HB pacing: premature extra-stimuli are delivered after a drive train of 600 ms at progressively shorter coupling intervals. Blue and burgundy circles denote the sudden change of QRS morphology from ns-HB QRS morphology to RV-myocardial QRS morphology. At coupling intervals of 450 – 340 ms, only ns-HB QRS morphology is present (HB effective refractory period of 330 ms). At coupling intervals of 330-250 ms, only RV-myocardial QRS morphology is present (RV myocardium effective refractory period of 250 ms). During relative refractory period of the HB (340-330 ms), there is already some QRS prolongation due to the HV interval prolongation and hence, a smaller contribution of the HB depolarization wavefront to the fused ns-HB QRS complex. During the relative refractory period of the RV myocardium (270-250ms), some QRS widening can also be observed. An ECG of the same patient is also presented on Supplementary Figure 1 – documenting identical QRS morphology during loss of HB capture with differential pacing output method. The blue bar corresponds to HB / RV capture (excitable period), the dashed bar to capture with decremental conduction (relative refractory period), and the burgundy bar to loss of capture (effective refractory period).

**Supplementary Figure 2.**
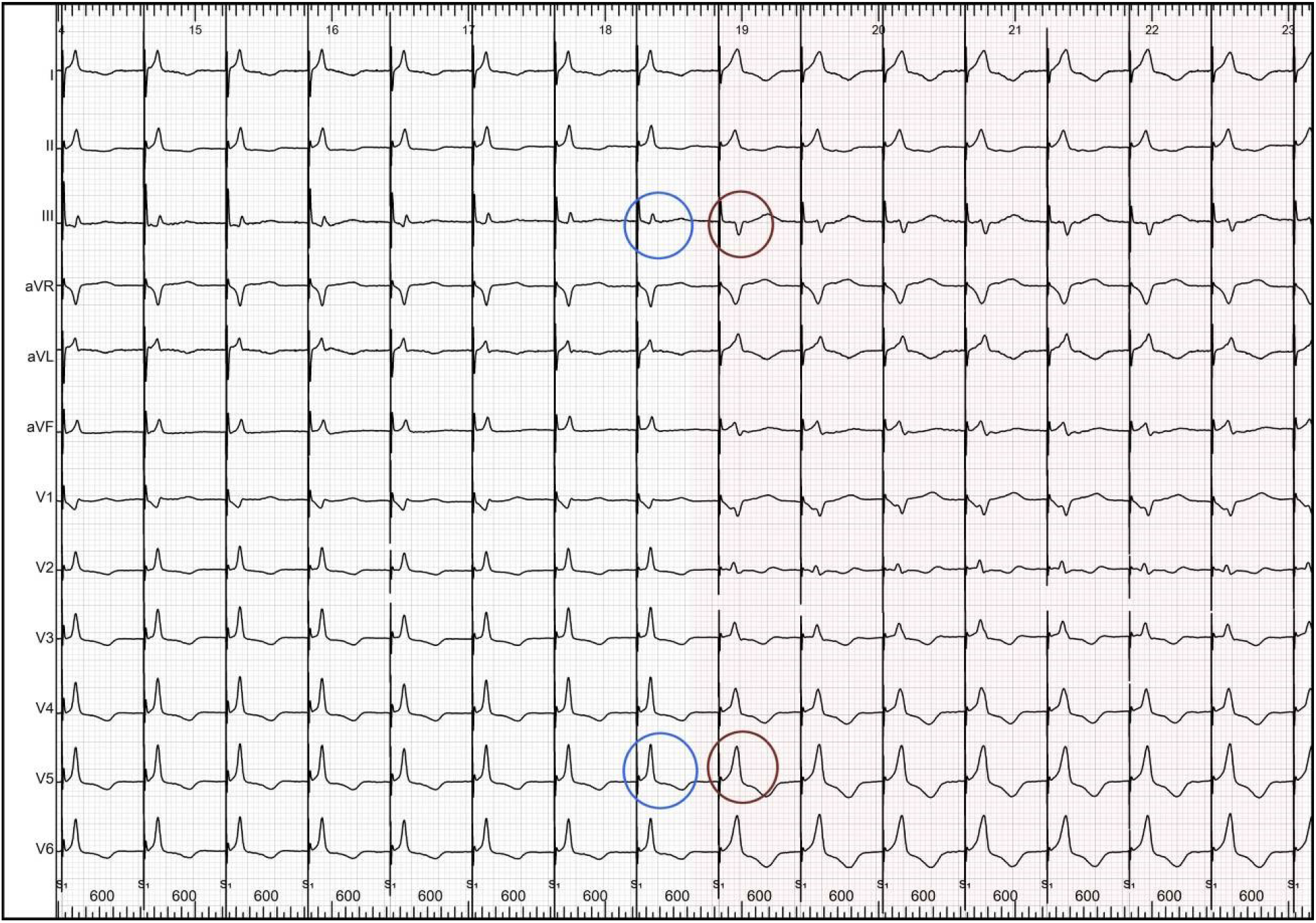
A 12-lead ECG illustrating the change of ns-HB QRS morphology into RV-myocardial QRS morphology when HB capture is lost during the lowering of the pacing output (differential pacing output method). Circles mark transitions from ns-HB QRS into RV-myocardial QRS; note the change of polarity in lead III, prolongation of QRS and less spiky R wave peak in I and V4-V6.

**Supplementary Figure 3.**
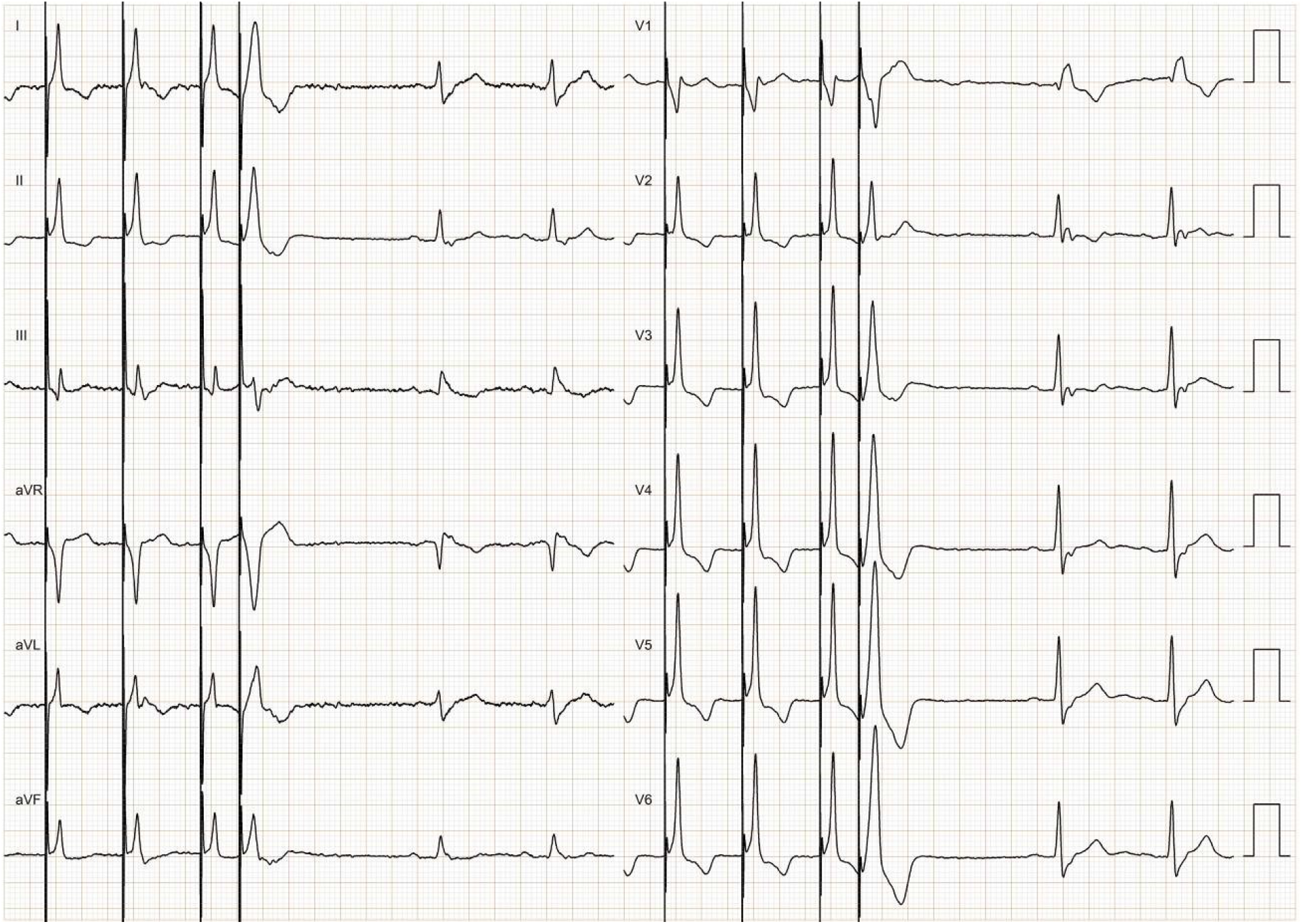
Programmed His Bundle pacing: a single extra-stimulus at the coupling rate of 300 ms reveals RV-myocardial capture QRS morphology (note, QRS prolongation, appearance of a notch in V1, rounding of R wave peak in I and change of polarity in lead III). At a coupling interval of 300 ms HB, is nearly always already refractory while the RV myocardium is still not. Thus, in the majority of cases, a response diagnostic of His bundle capture during non-selective pacing can be obtained with the introduction of an extra-stimulus at a single coupling interval.

